# Influence of ocean warming and acidification on juveniles of the true giant clam, *Tridacna gigas,* and its microalgal symbionts

**DOI:** 10.64898/2026.03.11.711206

**Authors:** Jake Ivan P. Baquiran, Niño Posadas, Michael Angelou L. Nada, Gabriella Juliane L. Maala, Patrick C. Cabaitan, Cecilia Conaco

**Author notes:** Corresponding author: Cecilia Conaco.

## Abstract

Uncontrolled carbon dioxide emissions from human activities contribute to ocean warming and acidification. These alterations in ocean chemistry threaten marine organisms, such as the true giant clam, *Tridacna gigas,* which is already imperiled due to overharvesting and habitat destruction. To gain an understanding of the physiological and molecular responses of *T. gigas* and its symbiotic dinoflagellates to ocean warming and acidification, we subjected juvenile individuals to different treatments simulating predicted seawater pH (7.6 and 8.0) and temperature (28°C, 30°C, 32°C and 34°C) levels for the next century. Juvenile giant clams were able to tolerate sustained exposure to temperatures of up to 32°C and pH as low as 7.6, while exposure to higher temperature (34°C), regardless of pH level, resulted in total mortality after a week. However, symbiosis was compromised even in the sublethal treatments, as indicated by the decrease in Symbiodiniaceae density and changes in symbiont gene expression. Symbionts significantly upregulated genes involved in splicing, translation, fatty acid metabolism, and DNA repair, which may constitute an adaptive response, while downregulating genes involved in photosynthesis and transmembrane transport, suggests impaired transfer of photosynthates to the host. These findings demonstrate the vulnerability of the juvenile *T. gigas* holobiont to heat stress, highlighting the critical importance of continued conservation and management alongside efforts to mitigate global changes in ocean conditions to safeguard this iconic marine bivalve.

**Summary Statement:** This study investigates physiological and molecular responses of *Tridacna gigas* to seawater warming and acidification, providing insights into the potential future of endangered giant clam populations in a changing ocean.

## Introduction

The ocean serves as a major sink of carbon dioxide (CO_2_), absorbing about 26% of the total anthropogenic greenhouse gas emissions (Mikaloff-Fletcher et al., 2006; Le Quere et al., 2014; Friedlingstein et al., 2019). Excessive amounts of dissolved CO_2_ alters the seawater carbonate chemistry and results in gradual decrease of pH or ocean acidification (OA). Due to positive radiative forcing brought by increased atmospheric CO_2_, ocean warming (OW) is predicted to co-occur with OA (Hoegh-Guldberg et al., 2007). The Intergovernmental Panel on Climate Change (IPCC) predicts that seawater pH will decrease by 0.3-0.5 units, while sea surface temperature may increase by ∼1.5-4.0°C or more by 2100 (Caldeira and Wickett, 2005; IPCC, 2014; Almazroui et al., 2020; IPCC, 2023).

Various studies have demonstrated the detrimental impacts of isolated or combined effects of ocean warming and acidification on diverse marine biota (Brennand et al., 2010; Navarro et al., 2016; Espinel-Velasco et al., 2018, Posadas et al., 2021, Giddens et al., 2022). Studies revealed that marine calcifiers, such as bivalve mollusks, are susceptible to future ocean conditions (Fujii et al., 2023; Mele et al., 2023). Acidified seawater can reduce calcification and promote the dissolution of CaCO_3_ structures (Leung et al., 2022). On the other hand, ocean warming impacts the soft tissue components and organs, inducing oxidative stress, altering energy metabolism, and causing structural damage (Masanja et al., 2023; Mao et al., 2025).

Giant clams (Tridacninae) are the largest and most conspicuous of the bivalves. There are 12 extant species: *Hippopus porcellanus, H. Hippopus*, *Tridacna squamosa*, *T. noae*, *T. maxima*, *T. crocea*, *T. derasa*, *T. mbalavuana, T. squamosina, T. rosewateri, T. elongatissima,* and *T. gigas* (Tan et al., 2021) distributed throughout the Indo-Pacific (Mies, 2019; Tan et al., 2022). Giant clams host ectosymbiotic photosynthetic dinoflagellate algae from family Symbiodiniaceae (Trench et al., 1981) in zooxanthellal tubules that originate from the gut and branch out in the colorful outer mantle (Norton et al., 1992; Hernawan, 2008). Algal symbionts contribute to the growth and nutrition of the giant clam host by transferring photosynthates, while benefiting from protection and host-derived nutrients (e.g. nitrogenous waste) (Hernawan, 2008; Soo and Todd, 2014; Neo et al., 2015; Logan et al., 2021). However, giant clams can bleach and lose their symbionts under stressful conditions (Sayco and Kurihara, 2024; Maboloc et al., 2025).

Among the tridacnids, the true giant clam, *T. gigas,* is the largest, growing up to 1.3 m in length and weighing more than 500 kg (Lucas, 2014). This species is classified as “critically endangered” by the International Union for Conservation of Nature (IUCN) Red List of Threatened Species, as it faces high risk of extinction (Wells, 1996; Neo and Li, 2024), mainly due to overharvesting for shell and meat trade (Mingoa-Licuanan and Gomez, 2002). In the Philippines, wild *T. gigas* are already considered locally extinct (Mingoa-Licuanan and Gomez, 2002; Cabaitan and Conaco, 2017).

Aside from overexploitation, giant clams are also susceptible to global change due to ocean acidification and warming (Watson and Neo, 2021). Individually, these factors influence survivorship, growth, reproduction, and calcification of giant clam species. In adult *T. crocea*, elevated temperature resulted in a pronounced reduction in survival rates and growth performance (Lee et al., 2024). Heat stress-induced bleaching also reduced the reproductive output of *T. crocea* by disrupting gametogenesis through egg resorption (Sayco et al., 2024). In addition, elevated levels of CO_2_ reduced the survival and growth of juvenile *T. squamosa* (Watson, 2015) and reduced growth in juvenile *T. crocea* (Kurihara and Shikota, 2018). In *T. maxima,* the holobiont thermal stress response was accompanied by differential regulation of genes involved in maintaining redox homeostasis (e.g., reactive oxygen species metabolic processes and oxidation-reduction processes), as well as some components of the symbiont photosynthetic machinery (e.g., photosystem II) (Alves Monteiro et al., 2020). At high temperature, *T. crocea* expressed genes involved in oxidative stress response, activation of apoptosis, and disruption of symbiosis in its outer mantle (Zhou et al., 2019). Ocean acidification, on the other hand, did not cause a significant reduction in survival and shell growth in juvenile *T. squamosa,* but transcriptome analysis revealed inhibition of calcification processes, as well as metabolic depression (Li et al., 2022). Whether warming and acidification induce synergistic or antagonistic responses, and whether these responses vary across giant clam species, remains to be understood.

Elucidating the molecular mechanisms underlying the response of the giant clam host and its algal symbionts to stressors can provide insights into the adaptive potential of *T. gigas*. Here, we subjected *T. gigas* juveniles to different levels of warming and acidification. We observed that temperature had a greater effect on survival and symbiont density in the giant clams. Notably, warming by itself or in combination with acidification induced global changes in gene expression, especially in the symbionts. This work suggests that the endangered giant clam *T. gigas* is likely to be negatively impacted by predicted changes in ocean conditions due to the disrupted host-algal symbiosis. The information derived from this study provides a glimpse into the possible fate of giant clams in the future ocean, which may help inform future strategies to improve giant clam conservation and management efforts.

## Materials and methods

### Giant clam collection and experimental setup

One hundred twenty-eight (128) hatchery-bred *T. gigas* juveniles were obtained from the giant clam ocean nursery of the Bolinao Marine Laboratory (BML) in Silaqui Island (16°26.806′N, 119°55.352′E), Bolinao, Pangasinan, northwestern Philippines. The giant clams were 2 years old with a mean shell length of 9.79 ± 0.05 cm (mean ± standard error) and a mean wet weight of 131.16 ± 2.35 g. They were cleaned of epibionts and parasites (e.g., pyramidellid snails) and were acclimatized for three days in a flow-through tank at the land-based hatchery of BML prior to the experiment.

The experiment was conducted using a flow-through ocean acidification and warming simulation system (Fig. S1). Sand-filtered seawater was cooled to ∼28°C using a 1HP recirculating chiller (HC-1000A, Hailea) then pumped into eight (8) 40-L rectangular plastic tanks or sumps where seawater pH was kept either at 8.0 by bubbling aeration, or adjusted to 7.6 using a mass flow controller unit connected to an air compressor (Hitachi, Japan) and carbon dioxide tank with pressure regulator (Yamato, Japan). A total of eight (8) sumps (4 at each pH setting) were used to supply seawater to 10-L glass aquaria at a flow rate of 5.4-7.2 L/hr. Aquaria were placed inside 40-L tanks with flowing seawater that could be adjusted to the target temperatures of 28°C, 30°C, 32°C, and 34°C using 300W thermostat heaters (Eheim, Germany). Additional flow in the tanks was provided by 200 L/hr submersible pumps (Hailea HX-1000). Tanks were illuminated with LED lamps on a 12 h light:12 h dark cycle.

### Ocean warming and acidification simulation experiments

Eight (8) treatments with different combinations of pH and temperature were tested: Present Day control (T1: 28°C and pH 8.0); Warming (T2: 30°C and pH 8.0; T3: 32°C and pH 8.0; T4: 34°C and pH 8.0); Acidification (T5: 28°C and pH 7.6); and Combined Warming and Acidification (T6: 30°C and pH 7.6; T7: 32°C and pH 7.6; T8: 34°C and pH 7.6). Treatments were selected to simulate predicted 2100 Representative Concentration Pathway (RCP) 2.6, 3.5, 8.5 scenarios (IPCC, 2023). The control (T1) was based on the yearly average seawater pH and temperature at the BML giant clam ocean nursery. Each treatment was represented by four (4) replicate aquaria that were placed in different 40L tanks so that no treatment replicates were dependent on the same sump. Four juvenile *T. gigas* individuals were randomly distributed into each of 32 glass aquaria.

Before the start of the experiment, all tanks were set at 28°C and pH 8.0. Juveniles were allowed to acclimatize in the aquaria for 5 days. After this period, the conditions were gradually adjusted over 6 days (+1.0°C and −1.0 pH unit per day) until the target temperatures and pH levels were reached. Juveniles were maintained at the desired conditions for 7 days (Fig. 1A-B). The experiment ran from 29 September 2018 (day 0) to 17 October 2018 (day 18) and was terminated when 100% mortality was observed in the most extreme treatments. All aquaria were cleaned every other day to minimize fouling.

**Fig 1.**
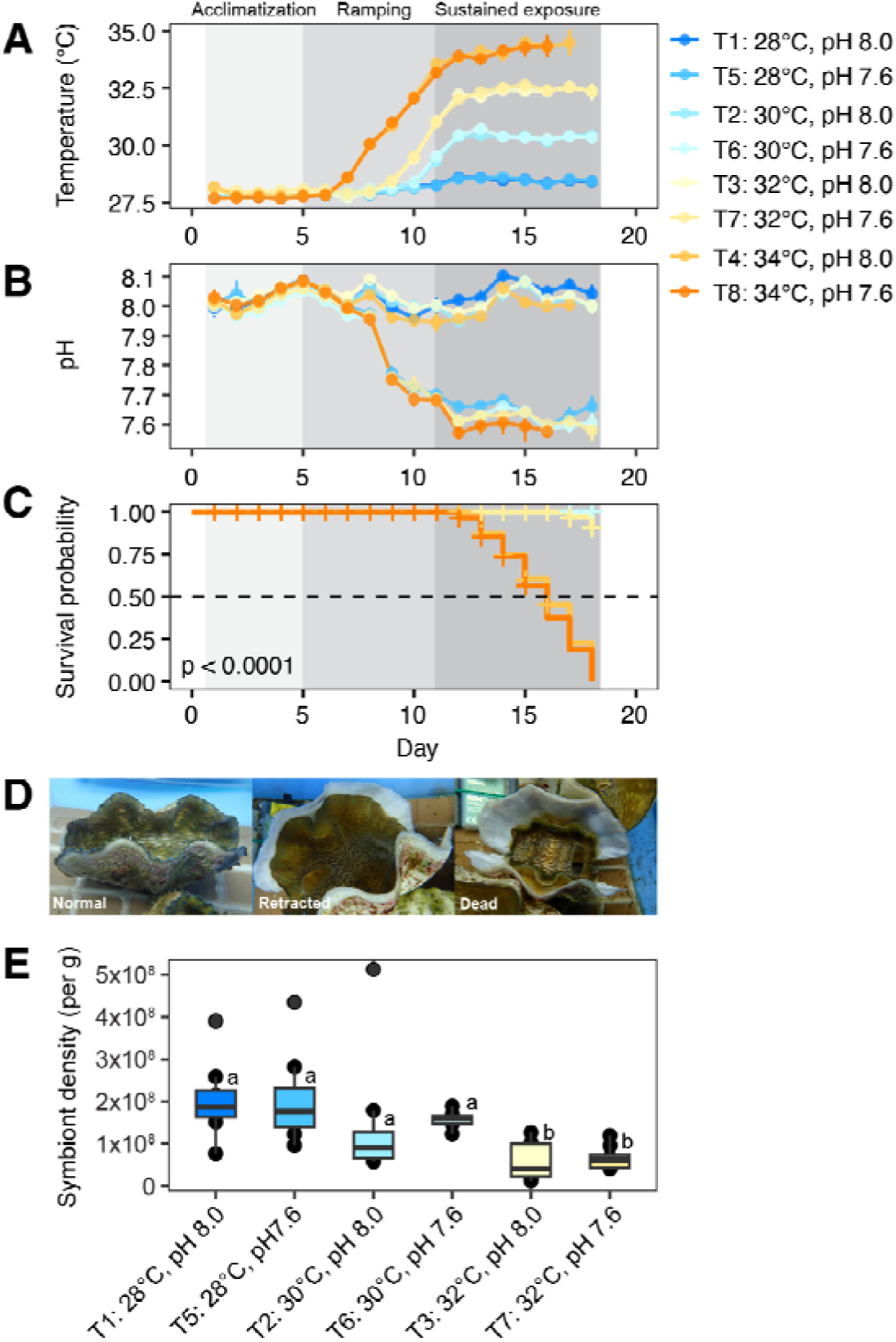
Effects of warming and acidification on juvenile *Tridacna gigas*. (A) Temperature profile, (B) pH profile, and (C) survival probability over the course of the experiment. Shaded areas in A-C indicate periods of acclimatization, ramping, and sustained exposure. (D) Representative images of normal, retracted, and dead giant clam juveniles. (E) Symbiont density in the mantle of surviving giant clams on day 18. Letters indicate statistically significant differences.

### Monitoring of seawater parameters

Physicochemical parameters in all aquaria were regularly monitored. Light intensity was recorded throughout the experiment using HOBO pendant data loggers (Onset Computer Corp., Bourne, MA, USA) positioned in each aquarium. Seawater pH was monitored using a portable pH meter (SG78 Mettler Toledo). Temperature, salinity, and dissolved oxygen (DO) were measured using a multi-parameter meter (YSI Pro2030). Water samples for total alkalinity (TA) were taken on days 3 and 6. TA was measured using a TA Analyzer (Kimoto Electric, ATT-05, Japan) calibrated using certified reference material (CRM) for ocean CO_2_ measurements. The co2sys.xls program (Pelletier et al., 2007) was used to calculate seawater *p*CO2, forms of inorganic carbon (e.g. bicarbonate ion (HCO_3_^-^), carbonate ion (CO_3_^2-^), and other carbonate system parameters such as aragonite (Ω_ara_) and calcite saturation (Ω_cal_) in the aquaria using the dissociation constants K1 and K2 from Mehrbach et al. (1973). Missing input values were replaced by the default values specified in the program. Seawater conditions remained relatively stable once the set points were reached (Table S1).

### Assessment of survivorship and symbiont density

Mortality of *T. gigas* individuals was recorded daily. Giant clams exhibiting degradation of the mantle or that had mantles completely separated from the shell were considered dead and were removed as soon as they were observed to maintain good seawater quality. At the end of the experiment (day 18), approximately 0.25 cm^2^ of the outer mantle of surviving clams (n = 8 per treatment) were non-lethally excised using surgical scissors and forceps for symbiont quantitation. The tissues were weighed then homogenized in 15 ml UV filtered seawater (UVFSW) using a tissue homogenizer (OMNI TH220) with disposable rotor stator generator probes. The homogenate was serially filtered through 120 µm, 60 µm, and 25 µm mesh sieves before centrifugation. Symbiont pellets were resuspended in filtered seawater to a final volume of 30 ml. Cell density was counted using a hemocytometer (Neubauer, Marienfeld, Germany). Values were expressed as cells per gram of tissue. Only intact algal cells with no visible signs of deformation were counted. No symbiont measurements were obtained from clams in T4 and T8 setups due to mass mortality.

### RNA isolation

Mantle clips for RNA isolation were taken at day 18, as described above. Tissues were flash frozen in liquid nitrogen then stored at −80°C. Tissues were collected from 4 individuals from treatments T1, T3, T5, and T7, one from each replicate tank (n = 4 per treatment). Giant clams from treatments T2 and T6 showed similar survivorship and symbiont density responses as individuals in treatments T1 and T5, and were not included in metatranscriptome analysis. Total RNA was extracted from frozen mantle tissues using TRIzol (Invitrogen) following the manufacturer’s recommendations. Contaminating genomic DNA was removed using the DNA-free kit (Invitrogen) and nucleic acid integrity was checked through agarose gel electrophoresis.

### Metatranscriptome sequencing, assembly, binning, and annotation

High-quality RNA samples were sent to Macrogen Inc., South Korea, for library preparation using the TruSeq Stranded mRNA LT Sample Prep Kit (Illumina, Inc., San Diego, CA, USA). Libraries were prepared from four samples per treatment representing different giant clam individuals. mRNA-enriched libraries were sequenced on the Illumina NovaSeq6000 platform to generate 100 bp paired-end reads. Raw reads were quality checked using FastQC v.0.11.8 (https://www.bioinformatics.babraham.ac.uk/projects/fastqc/) and trimmed using Trimmomatic v.0.39 (Bolger et al., 2014). Adapter sequences were removed, poor-quality bases (quality score below 3) at leading and trailing bases were trimmed, and reads less than 50 bases long were eliminated. *De novo* transcriptome assembly was carried out using Trinity v.2.8.5 (Grabherr et al., 2011). Transcripts with 90% similarity were clustered and the longest representative contigs retained using CD-HIT-EST v.4.8.1 (Fu et al., 2012) then reads were mapped back to the clustered assembly. Transcripts shorter than 300□ bp and isoforms with zero isoform percentage (IsoPct) were removed to eliminate putative fragmented and misassembled transcripts. Peptide sequences encoded by the longest open reading frame for each transcript were predicted using TransDecoder v. 5.5.0. Raw sequence reads were deposited in the NCBI Short Read Archive database under BioProject PRJNA1430147. The non-redundant reference transcriptome is available at FigShare (https://doi.org/10.6084/m9.figshare.31442209). Predicted peptide sequences were annotated by Blastp (e-value < 1x10^-5^) using DIAMOND (v.0.9.24) (Buchfink et al., 2014) against the SwissProt, and NCBI non-redundant (nr) protein databases. Gene ontology annotations were obtained using Blast2GO (Conesa et al., 2005) with the top SwissProt hit for each peptide as input. A comprehensive annotation report was generated using Trinotate.

Host and symbiont transcripts were identified based on taxon affiliations of best hits to the NCBI nr protein database. Transcripts with best hits to Metazoa were classified as host transcripts and those with best hits to Alveolata were considered symbiont transcripts. Transcripts aligning to other taxa or with no matches in the nr database were excluded from further analyses. Completeness of the host and symbiont bins was assessed by comparing against the Mollusca and Alveolata reference databases (odb_12) in BUSCO version 6.0.0. (Simao et al., 2015). Host-assigned transcripts were also aligned against the predicted proteins in the *T. gigas* genome (Li et al., 2024). Symbiont sequences were aligned to genomes of other Symbiodiniaceae representatives, including *Symbiodinium microadriaticum* (Aranda et al., 2016), *Breviolum minutum* (Shoguchi et al., 2013), *Cladocopium infistulum* (González-Pech et al., 2024), *C. proliferum* (Liu et al., 2018), *Cladocopium* sp. (Y103 isolate) (Shoguchi et al., 2018), *Cladocopium* sp. (C15 ITS2-subtype) (Robbins et al., 2019), *Durusdinium trenchii* (Shoguchi et al., 2021), and *Fugacium kawagutii* (Liu et al., 2018) using Blastp (e-value < 1 × 10^−5^). Symbiont affiliations of transcripts were assigned based on top Blastp hit (highest % identity and lowest *e*-value).

### Differential gene expression analysis

Trimmed reads were mapped back to the transcriptome assembly using RSEM (Li and Dewey, 2011). Differential gene expression analysis was done using the Wald test in DESeq2 (Love et al., 2014) to compare gene expression in T3, T5, and T7 relative to the control (T1). Only genes with > 1 count in at least four libraries were included in the analysis. Genes with an FDR-adjusted p-value < 0.05 were considered differentially expressed. Genes with differential isoform usage in the treatments were identified using Iso-maSigPro in the maSigPro R package (Nueda et al., 2017).

### Gene ontology and pathway enrichment analysis

Functional enrichment analysis was conducted using the topGO package in R (Alexa and Rahnenfuhrer, 2009). GO terms with an FDR-adjusted p-value < 0.05 were considered significantly enriched. Significantly enriched biological processes (BP) were summarized using the rrvgo package in R (Sayols et al. 2023).

Predicted peptides of symbionts were aligned against the STRING v.11 (Mering et al., 2003) protein database for *Arabidopsis thaliana,* using an e-value cut-off of 1x10^-5^. Interaction networks for homologous proteins in STRING (confidence > 0.7) were visualized using Cytoscape v.3.10.4 (Shannon et al., 2003).

### Statistics

Survival of *T. gigas* in the different treatments was visualized using Kaplan-Meier survival curves and the statistical difference was computed using the survfit function. Differences in symbiont density were determined using the Kruskal-Wallis test with Wilcoxon test for multiple pairwise comparisons. All statistical analyses and data visualization were done on R v.4.5.1 (R Development Core Team, 2010) using packages Survminer (Kassambara et al., 2021), Ryan Miscellaneous (Rmisc) (Hope, 2013), vegan (Oksanen et al., 2017), Simple Fisheries Stock Assessment Methods (FSA) (Ogle, 2017), and ggplot2 (Wickham, 2016).

## Results

### Effects of lowered pH and elevated temperature on the T. gigas holobiont

Juvenile *T. gigas* showed 100% survival over the 18-day duration of the experiment in treatments with temperatures set at 28°C or 30°C regardless of pH level (T1, T5, T2, T6; Fig. 1C, Dataset 1). Juveniles in treatments set at 32°C (T3, T7) exhibited some mortality starting at day 17 (after 5 days of sustained exposure), but individuals did not exhibit any mantle retraction or visible paling, and survival rate was at 94% by the end of the experiment. In contrast, giant clams in treatments set at 34°C (T4, T8), at either pH level, exhibited severe mantle retraction and mortalities starting on day 12 (Fig. 1D), with no survivors by the end of the experiment. Mantle symbiont density on day 18 was 2.05 ± 0.32 x 10^8^ cells per g of tissue (mean ± standard error) in the control (T1). A similar density of 2.06 ± 0.38 x 10^8^ cells per g was recorded in the acidified treatment at ambient temperature (T5). In the treatments set at 30°C, symbiont densities ranged from 1.46 ± 0.54 x 10^8^ cells per g in T2 and 1.58 ± 0.06 x 10^8^ cells per g in T6 (Fig. 1E; Table S2). Giant clams in treatments set at 32°C showed significantly lower symbiont density (p < 0.05), with 5.78 ± 0.15 x 10^7^ cells per g in T3 and 6.65 ± 0.99 x 10^7^ cells per g in T7.

### De novo metatranscriptome assembly

RNA sequencing yielded 616,614,396 paired-end Illumina reads from 16 juvenile *T. gigas* samples from four treatments. Filtering of the *de novo* transcriptome assembly resulted in a final non-redundant assembly of 46,296 protein-coding genes (54,170 transcripts), of which 16,837 were from the host and 29,459 were from the symbionts (Table S3). Of the host-assigned genes, 95% (15,983 genes) aligned to sequences in the *T. gigas* genome. The host gene set represented 86.7% of the BUSCO molluscan database, while the symbiont gene set represented 94.9% of the BUSCO alveolate database. Majority (90%) of symbiont transcripts aligned best to the genomes of *Cladocopium,* indicating that the dominant symbiont of *T. gigas* belongs to this genus (Table S4).

### Effects of lowered pH and elevated temperature on gene expression

A total of 2,933 genes were differentially expressed (FDR-adjusted p-value < 0.05) in the acidification (T5: 28°C, pH 7.6), warming (T3: 32°C, pH 8.0), and combined warming and acidification (T7: 32°C, pH 7.6) treatments relative to Present Day controls (T1: 28°C, pH 8.0) (Dataset 2). In the acidification treatment, only 3 genes were differentially expressed, all of which were from the symbionts. In the warming treatment, a total of 542 genes were differentially expressed (4 upregulated and 26 downregulated in the host; 354 upregulated and 158 downregulated in the symbionts). In the combined treatment, a total of 2,855 genes were differentially expressed (30 upregulated and 45 downregulated in the host; 1,570 upregulated and 1,210 downregulated in the symbionts). Majority of the differentially expressed transcripts were of symbiont origin (Fig. 2A). Only one differentially expressed gene was common among all treatments and 465 were common in the warming and combined warming and acidification treatments (Fig. 2B).

**Fig 2.**
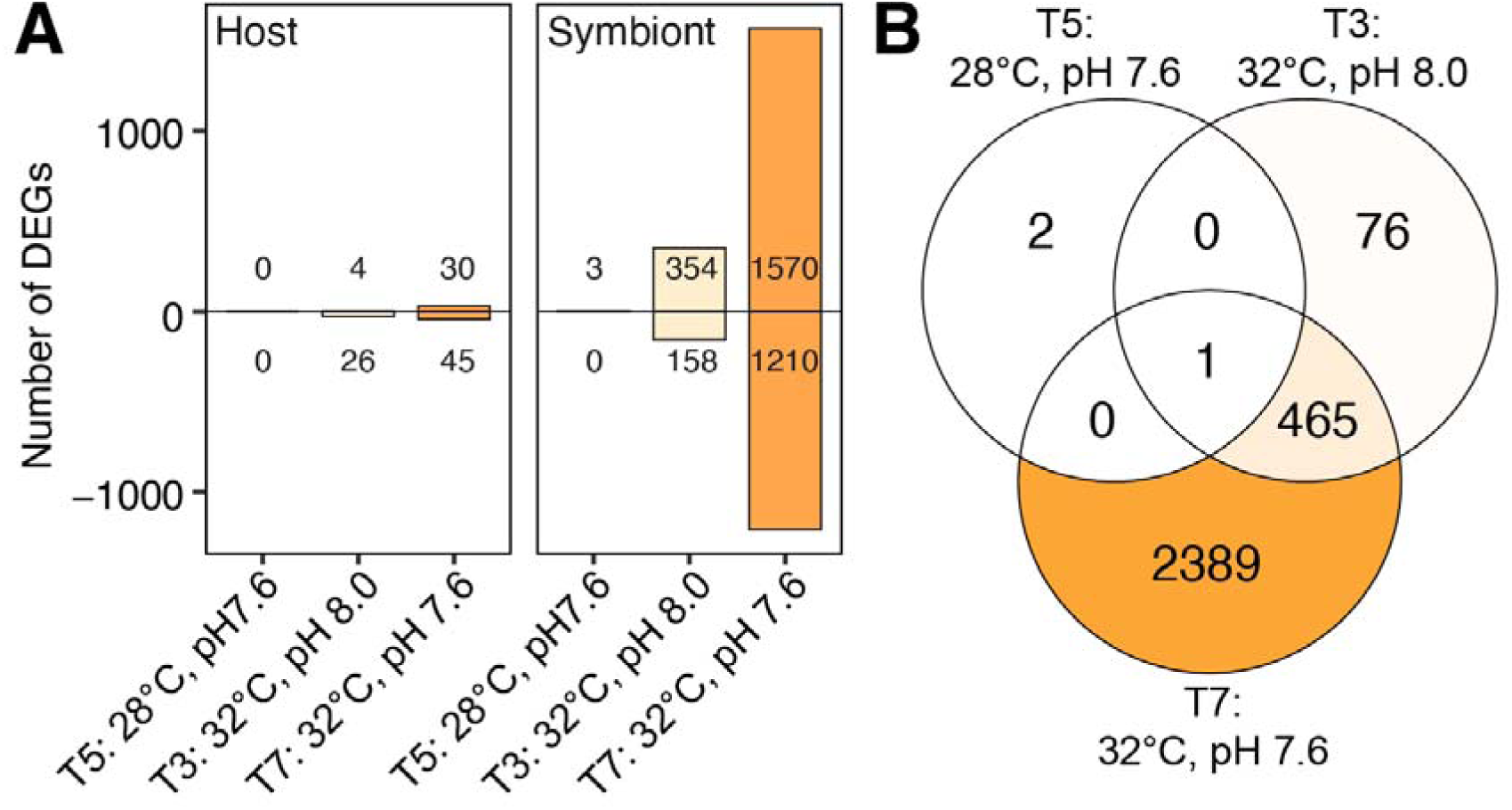
Differential gene expression in *Tridacna gigas* under warming and acidification. (A) Number of differentially expressed genes (FDR-adjusted p-value < 0.05) in host and symbionts in the acidification (T5: 28°C, pH 7.6), warming (T3: 32°C, pH 8.0), and combination (T7: 32°C, pH 7.6) treatments. (B) Number of differentially expressed genes that are common or unique across treatments.

### Differential gene expression in the giant clam holobiont

Acidification, warming, and combination treatments elicited a minimal shift in transcriptional profile in the giant clam host. No differentially expressed genes were detected under acidification alone. In the warming and combined warming and acidification treatments, genes related to DNA replication (e.g. helicase MOV10, topoisomerase TOP3B, sentrin protease SENP7, ligase DNLI3), heat shock proteins (HS71L), and E3 ubiquitin ligases (RCBT1 and DZIP3) were upregulated. Genes related to defense (e.g. lectins CLC4M and COL10, cytochrome P450, tyrosine-protein phosphatase PTP), fatty acid uptake and metabolism (FABP6), ion channel conductance (CLC4A), RNA processing (HRB87F, HNRNPA1), and muscle control (octopine dehydrogenase OCDH, tubulin polyglutamylase TTLL6, MOXD2) were downregulated (Dataset 2).

In contrast to the host, giant clam symbionts exhibited a robust transcriptional response, particularly in the warming and combination treatments. Functions overrepresented in the upregulated gene set included translation, mRNA splicing, DNA damage response, fatty acid metabolism, amino acid metabolism, and response to heat (Fig. 3; Dataset 3-4). The downregulated gene set included functions related to photosynthesis light reaction, nitrate assimilation, isoprenoid synthesis, microRNA-mediated silencing, response to stress, ion homeostasis, protein import into the chloroplast, antibiotic metabolism, and transmembrane transport. The symbiont nucleic acid binding protein, FUBP3, was upregulated in all treatments, including acidification (Dataset 2).

**Fig 3.**
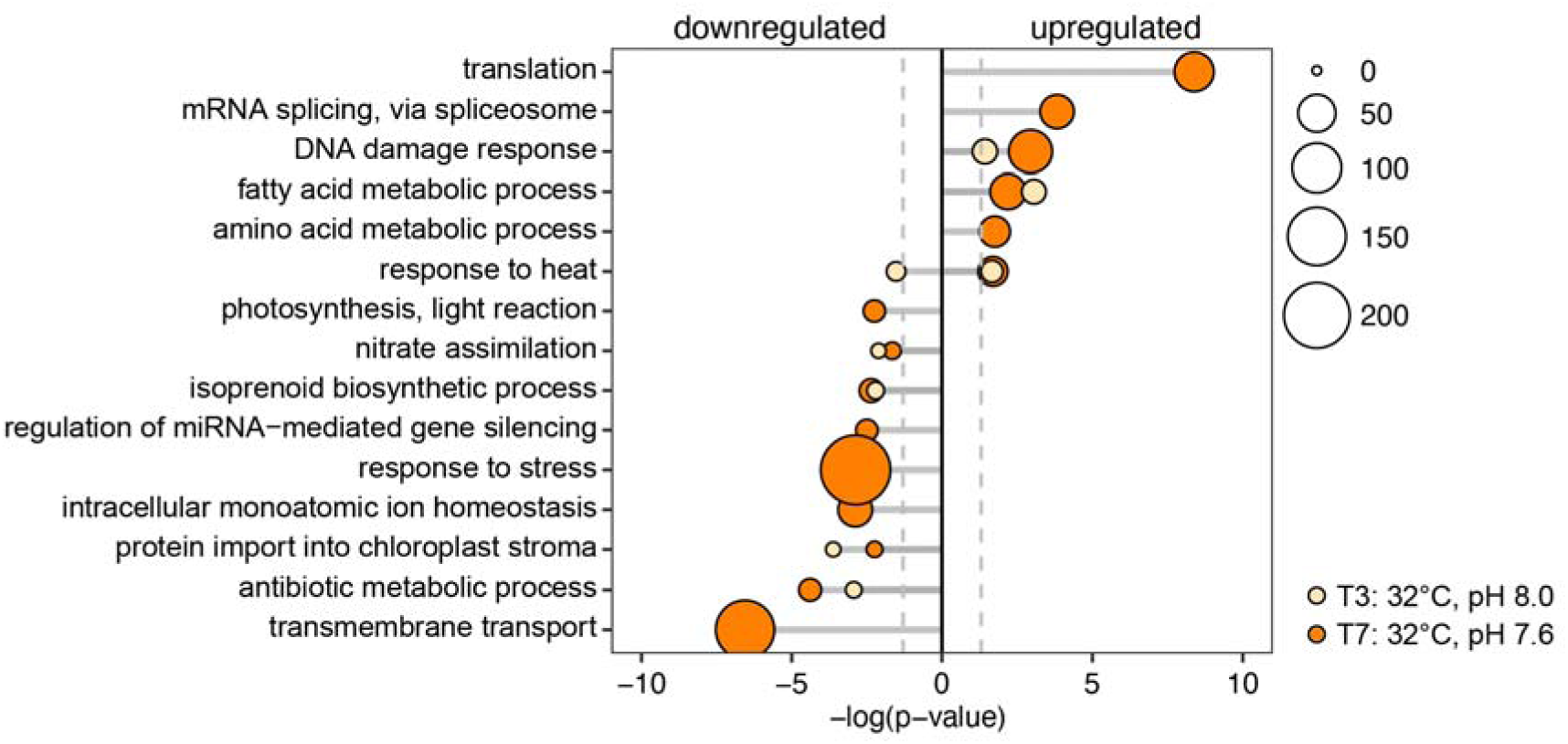
Symbiont gene functions affected by warming and acidification treatments. Selected functions enriched in differentially expressed (DE) genes in the giant clam symbionts in the warming (T3: 32°C, pH 8.0; yellow) and combination (T7: 32°C, pH 7.6; orange) treatments. Only terms with FDR-adjusted p-value < 0.05 across treatments were included. Dashed lines demarcate the significance threshold. Marker size corresponds to the number of differentially expressed genes in each category.

Homologs of differentially expressed genes in the giant clam symbionts under the combined warming and acidification treatment were mapped onto the *Arabidopsis* interaction network (Dataset 5). This revealed extensive interconnections among 543 genes representing enriched functions linked to metabolism, photosynthesis, stress response, and gene expression control (Fig. 4A). Genes involved in the spliceosome and DNA repair showed a greater proportion of upregulation (80-82%) compared to the other enriched functions (Fig. 4B).

**Fig 4.**
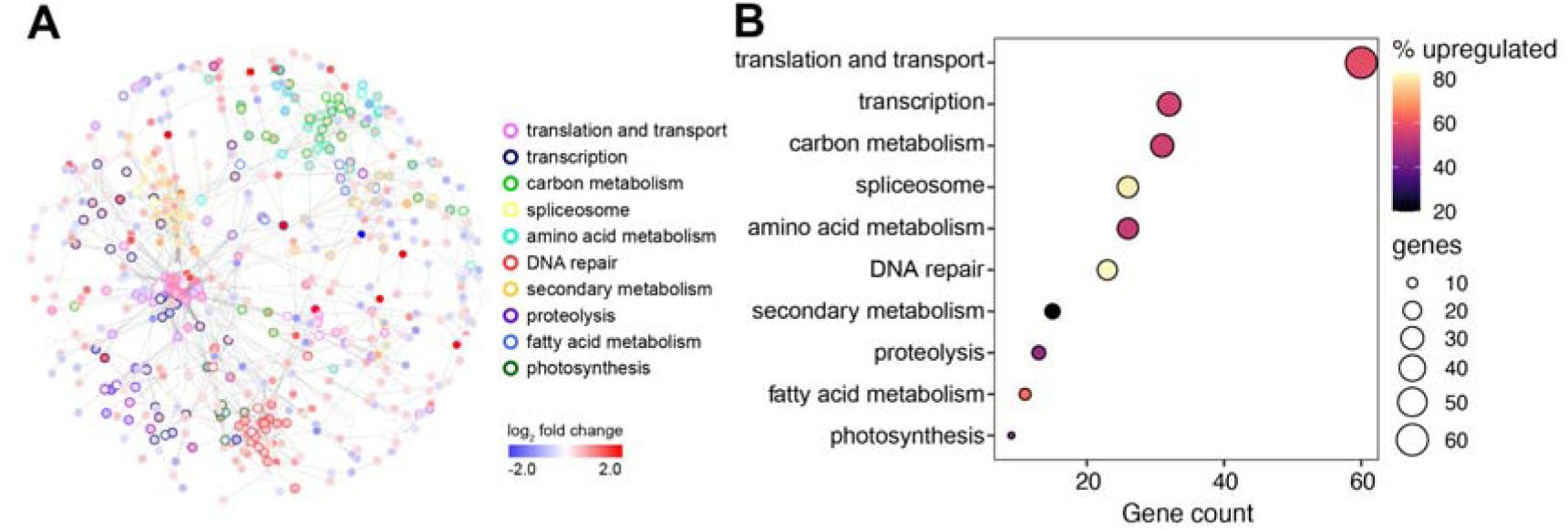
Gene networks affected by combined acidification and warming. (A) Interaction network of symbiont gene homologs. Node fill colors represent shifts in expression based on log_2_ fold change (red, upregulated; blue, downregulated) in the combined acidification and warming treatment (T7: 32°C, pH 7.6) relative to control. Selected functions are indicated by node border colors. The network is based on the *Arabidopsis* network in STRING. (B) Number of differentially expressed genes and the proportion that are upregulated in each functional category within the network.

Upregulation of multiple RNA splicing-related genes including splicing factors (CLO, U2AF65A, ATO, SC35), RNA helicases (RH20, RH21, RH45, EIF4A3, BRR2A), an RNA-binding protein (Q494N5), and other pre-mRNA-processing proteins (PRP40B and PRP19A) (Dataset 5) in the treatment prompted us to examine isoform switching in the giant clam symbionts. We discovered 23 genes with transcript isoforms that changed in expression across treatments. Among these were genes related to signaling (CACNA1), epigenetic control (JMJ8), metabolism (MNS1, MI2D, AATC), translation (AARS, DTWD2), protein folding (CALR), transport (KIF6, KIF9, ANTL2), and ubiquitination (FEM1B, UBC6) (Fig. S2; Dataset 6).

## Discussion

Stressors can significantly alter the physiology, behavior, and molecular processes of an organism depending on its tolerance threshold (Payton et al., 2016; Anestis et al., 2017). Here, we subjected *T. gigas* juveniles to different combinations of acidification and warming to assess physiological and transcriptional responses of the giant clam holobiont to predicted future ocean conditions.

### Thermal limits of Tridacna gigas

*Tridacna gigas* juveniles were capable of tolerating approximately 7 days of sustained exposure to acidification at −0.4 pH units from the control. These results are similar to reports in other bivalves, such as the grooved carpet clam, *Ruditapes decussatus* (−0.4 and −0.7 pH units; Range et al., 2011), and juvenile *Crassostrea virginica* and blue mussel, *Mytilus edulis* (−0.7 pH units; Stevens and Gobler, 2018), that can tolerate acidification with exposures ranging from 1-10 months. A few studies observed shell dissolution in bivalves subjected to acidified conditions (Mackenzie et al., 2014; Carlson et al., 2025). In the marine gastropod, *Lunella coronata granulata,* significant inhibition of shell growth was seen at pH 7.5 (−0.6 pH units) and pH 7.1 (−1.0 pH units), with intensified shell surface erosion and extensive periostracum peeling (Li et al., 2025).

Giant clam juveniles were also able to tolerate elevated temperature up to 32°C (+4°C from control). However, further raising the temperature to 34°C (+6°C from control) proved lethal to the organism. This level of temperature rise has previously been shown to be detrimental for survival of early developmental stages of giant clams, with 33°C (+5°C from control) reducing the survival rate of *T. gigas* larvae (Enricuso et al., 2018) and ∼29.5°C (+5°C from control) leading to complete mortality of *T. squamosa* trochophores after 24 hr (Neo et al., 2013). This delineates the upper limit of the “thermal window” (Bielen et al., 2016) for giant clam juveniles, above which their ability to maintain cellular homeostasis is severely compromised. Similar thermotolerance limits have been reported in other bivalves. For example, both *Scrobicularia plana* and *Cerastoderma edule* showed 100% mortality at 32°C (+9°C from optimum) after just 5 days of exposure (Verdelhos et al., 2015). Similarly, the manila clam, *Venerupis philippinarum,* showed ∼60% cumulative mortality after 8 days exposure to 24°C (+9°C from control) (Bae et al., 2021).

Giant clam survival in the treatments combining acidification and temperature stress was similar to the temperature-only treatments, suggesting that temperature was the main driver of the giant clam response. These findings mirror studies in the Eastern oyster, *Crassostrea virginica,* where warming (+3°C above ambient) but not acidification (ca. −0.2 pH units) reduced survival after around five months (Speights et al., 2017). Juvenile bay scallops, *Argopecten irradians,* also showed sensitivity to warming at 30°C (+6°C from ambient), irrespective of pH or dissolved oxygen level, with significantly decreased mortality after four weeks (Stevens and Gobler, 2018). However, other studies present contrasting results, with exacerbated effects from combined stressors. For example, the combination of seawater temperature at 31.5°C (+3°C from ambient) and 1019 µatm pCO_2_ or pH

∼7.8 (−0.3 pH units from control) resulted in only ∼20% survival of *T. squamosa* after 60 days of exposure (Watson et al., 2012). Elevated temperature of 28°C (+4°C from control) either with ambient or lower pH also decreased the survival of juvenile hard clam, *Mercenaria mercenaria,* and bay scallops, *A. irradians* (Talmage and Gobler, 2011). Variability in the responses of diverse bivalves to thermal and acidification stressors may reflect organismal physiology, but could also be influenced by differences in experimental conditions and exposure duration.

### Molecular responses of the giant clam host to warming and acidification

Gene expression responses in the giant clam holobiont varied across the treatments. Acidification by itself did not elicit much of a change in transcription, indicating that this condition and duration of exposure was not stressful to the organism. In contrast, warming triggered a bigger transcriptional response, which was further amplified in the combined acidification and warming treatment, demonstrating that combined stressors can have synergistic impact.

Although the *T. gigas* host did not exhibit a major shift in gene expression, significant downregulation of some defense-related genes suggests that the protective responses of the giant clam are attenuated. If the conditions are sustained over a longer period, it can be expected that this will further diminish the ability of the host to fend off opportunistic pathogens or disease-causing bacteria. This has been demonstrated in the mussels, *M. edulis* and *M. mercenaria*, which exhibit an increase in parasite and pathogen abundance under conditions of warming and acidification (Mackenzie et al., 2014; Schwaner et al., 2020). In addition, downregulation of genes related to cellular structure and muscle contraction may be linked to the weakening of giant clam tissues, which could contribute to mantle retraction under extreme or prolonged stress (Mao et al. 2025; Basti and Segawa, 2010).

### Warming and acidification disrupts host-algal symbiosis

Giant clams in the sublethal treatments (i.e. at temperatures below 34°C) did not exhibit obvious signs of stress, such as mantle retraction or bleaching. However, we did measure a significant decrease in symbiont density in the 32°C treatments. This suggests that symbiont expulsion from the host was already occurring and it is likely that mantle paling may become apparent if the exposure is extended past day 18. Other studies have observed declines in symbiont density in *T. maxima* exposed to 30.7°C (+1.5°C from ambient) for 65 days (Brahmi et al., 2021) and *T. squamosa* subjected to increased *p*CO2 (Li et al., 2022). Visible bleaching has also been reported in *T. crocea*, *T. squamosa, T. gigas,* and *T. maxima* subjected to 31-34°C (about +2-4°C from ambient) (Junchompoo et al., 2012; Sayco et al., 2023; Sayco et al., 2024; Teaniniuraitemoana et al., 2025).

Symbiont populations significantly decrease during bleaching, which disrupts the acquisition of photosynthates by the host (Leggat et al., 2003). When this happens, giant clams must rely on their ability for heterotrophic feeding. However, giant clams show varying levels of dependence on their symbionts, with *T. gigas* being most reliant on autotrophic sources of nutrition (Guibert et al., 2025). This giant clam species likely benefits from association with members of *Cladocopium* (Guibert et al., 2025), which are known to efficiently share photosynthates with their host (Stat et al., 2008; Poo et al., 2021). Thus, *T. gigas* is expected to be particularly vulnerable to starvation upon symbiont loss.

### Adaptive strategies of symbionts may compromise the holobiont

Transcriptional shifts observed in *T. gigas* juveniles was mostly due to the symbionts rather than the giant clam host. This is in stark contrast to corals where the host response is usually more robust because their intracellular symbionts are better insulated from external environmental fluctuations (Venn et al., 2025). In giant clams, symbionts are housed within zooxanthellal tubules in the mantle and may be more exposed to the environment, thus requiring them to respond dynamically to physicochemical changes.

Dinoflagellate symbionts of giant clams possess large and complex genomes with genes tandemly arranged in unidirectional clusters, requiring regulatory control at the post-transcriptional level (Janouskovec et al., 2017; Lin et al., 2024; Gavin et al., 2026). Some multi-gene transcripts expressed from gene arrays are trans-spliced with addition of a 5’ spliced leader in dinoflagellates (Zhang et al., 2007). Notably, the symbiont far-upstream element-binding protein 3 (FUBP3) was upregulated in all treatments. FUBP3 is a single-stranded DNA- and RNA-binding protein that has been implicated in various aspects of RNA metabolism, including splicing regulation (Li et al., 2013), and may play an important role in the symbiont gene expression response. Genes involved in cell signaling, including ion transport and ubiquitination, and environmental response, including nucleic acid metabolism, are also likely targets for splicing regulation in dinoflagellates (Song et al., 2017; Haro et al., 2025). The expression of alternative isoforms could allow symbionts to produce protein variants that may function more efficiently under prevailing conditions (Yang et al. 2022; Dougan et al. 2023). Further analysis of the functions of differentially expressed isoforms is needed to elucidate their roles in the symbionts under stress.

The processes of RNA splicing, translation, and fatty acid metabolism are active in giant clam symbionts subjected to warming and acidification. These may enable modification of cellular components, such as enzymes and lipid membranes, for enhanced performance in their new environment. In particular, changes in fatty acid metabolism have been reported in coral-associated *Symbiodinium* subjected to prolonged thermal stress (Gierz et al., 2017). Shifts in fatty acid and lipid profiles may support maintenance and repair of membrane structures at elevated temperature (Parkinson et al., 2016; Vidyarathna et al., 2024).

In contrast, photosynthetic activity of the symbionts may be inhibited by warming and acidification. This is supported by the reduction in light harvesting pigments, such as carotenoids and chlorophyll (Gong et al., 2021), as well as downregulation of genes for light reaction components, nitrate assimilation, and protein targeting to the chloroplast. Furthermore, although symbionts remaining in the giant clam mantle can still undertake photosynthesis, it is likely that translocation of photosynthetic products to the host is impaired, as evidenced by downregulation of transmembrane transport (Rädecker et al., 2021). Allocation of energy resources in the symbiont to processes like protein synthesis and fatty acid metabolism may result in a trade-off, with downregulation of functions related to photosymbiosis. Thus, while symbionts are working to maintain cellular homeostasis, decreased nutritional contribution to the host impairs symbiosis (Baker et al 2018; Allen-Walker et al., 2023), rendering the holobiont susceptible to sustained stress exposure.

The contrasting effects of warming and acidification stress on survival, symbiosis, and gene expression observed in *T. gigas* likely reflect a distinct mechanism of action of these stressors. While temperature can cause widespread protein denaturation and increased oxidative stress within a short period, organisms may be able to maintain intracellular pH homeostasis through the activity of proton pumps (Choi et al., 2021; Schwaner et al., 2022). In addition, rising temperature is typically associated with decreased levels of dissolved oxygen, which may contribute to mortality at higher temperatures. Simultaneous application of warming and acidification, on the other hand, may amplify the cellular response, resulting in far greater change in transcriptional profile than each stressor by itself, though in some cases this could overwhelm protective cellular mechanisms and deplete energy resources.

## Conclusion

The true giant clam, *T. gigas,* is now locally extinct in many areas within its range. Although efforts to protect existing individuals and to revitalize breeding populations continue, global stressors such as warming and acidification will affect giant clam survival at multiple stages of its life cycle. Understanding how the giant clam holobiont responds to predicted ocean conditions provides us with insights into how we may improve on existing conservation strategies, such as through selecting appropriate sites for restocking, enhancing habitat protection, provision of symbionts that best support the nutritional needs of the giant clam host, or even breeding for thermotolerance. The persistence of giant clams into the future will require a concerted effort, not only to prevent overexploitation, but also to mitigate the stressors that threaten their survival.

## Acknowledgements

We acknowledge the Bolinao Marine Laboratory for providing access to the Silaqui Giant Clam Ocean Nursery. We thank Francis Kenith Adolfo, Robert Valenzuela, Ben Jack Gabuay, and Aubrey Joy P. Tejada for assistance with field and laboratory experiments; Dr. Ma. Lourdes San Diego-McGlone and Natasha Tamayo for assistance with seawater analyses; Keana Dehnielle P. Tan, John Bennedick Quijano, and Kelly Rome A. Publico for assistance with data analysis. This study was funded by the Department of Science and Technology Philippine Council for Agriculture, Aquatic and Natural Resources Research and Development (QMSR-MRRD-MEC-295-1449) to PCC and CC.

## Author contributions

Jake Ivan P. Baquiran: Investigation, Methodology, Project administration, Formal analysis, Writing - original draft, Writing - review & editing

Niño Posadas: Formal analysis, Visualisation, Writing - original draft, Writing - review & editing

Michael Angelou L. Nada: Investigation, Methodology, Project administration, Formal analysis

Gabriella Juliane L. Maala: Formal analysis, Writing - review & editing

Patrick C. Cabaitan: Conceptualization, Fund acquisition, Writing - review & editing

Cecilia Conaco: Conceptualization, Fund acquisition, Methodology, Formal analysis, Supervision, Visualisation, Writing - original draft, Writing - review & editing

## Data availability

Raw sequence data used in this study are available for download at NCBI Short Read Archive database under BioProject PRJNA1430147.

The non-redundant transcriptome assembly, assembly annotation, and count data are available from Figshare at https://doi.org/10.6084/m9.figshare.31442209.

## Notes

https://doi.org/10.6084/m9.figshare.31442209

### Competing Interest Statement

The authors have declared no competing interest.

