## Supplementary information for "Influence of ocean warming and acidification on juveniles of the true giant clam, *Tridacna gigas,* and its microalgal symbionts"

*Corresponding author:

Cecilia Conaco

**ORCID**

Jake Ivan P. Baquiran https://orcid.org/0000-0001-7730-1061

Niño Posadas https://orcid.org/0000-0002-0583-5096

Michael Angelou L. Nada https://orcid.org/0000-0002-3466-3465

Gabriella Juliane L. Maala https://orcid.org/0009-0009-2955-289X

Patrick C. Cabaitan https://orcid.org/0000-0002-5207-2378

Cecilia Conaco https://orcid.org/0000-0002-3594-2810

**Supplementary information**

**Supplementary Figures**

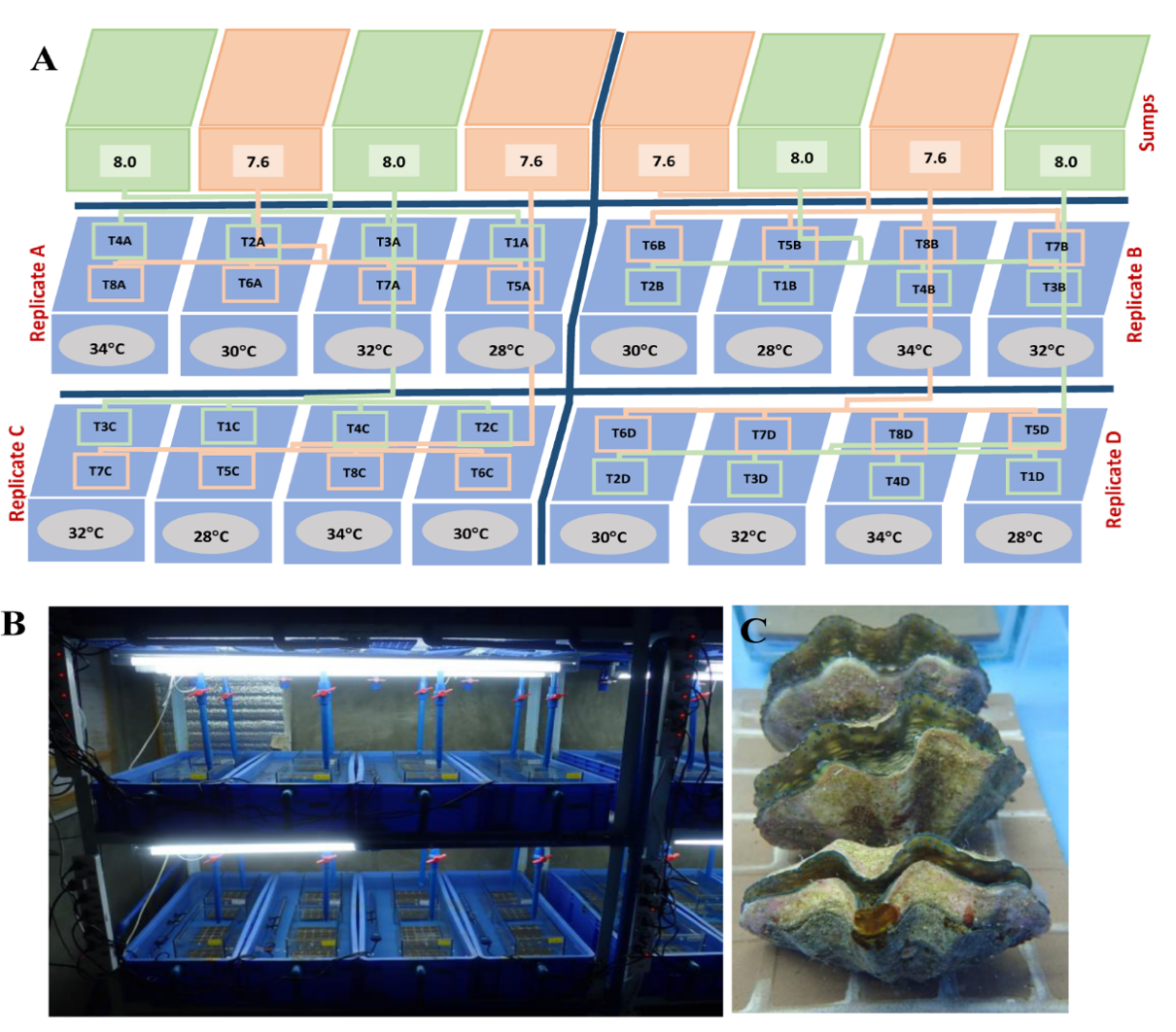

**Figure S1.** Experimental setup. (A) Design of the giant clam *ex situ* ocean acidification and warming experiment. Seawater pH was manipulated in 8 plastic tanks or sumps (40L; outer dimensions: 32 x 12 x 7 in). This seawater fed into glass aquaria (10L; outer dimensions: 12 x 8 x 10 in) placed inside 40L tanks (outer dimensions: 15 x 23 x 8 in) with flowing seawater that could be adjusted to the target temperatures using submersible heaters. (B) Photo of the setup. (C) Juvenile *T. gigas* individuals in a tank.

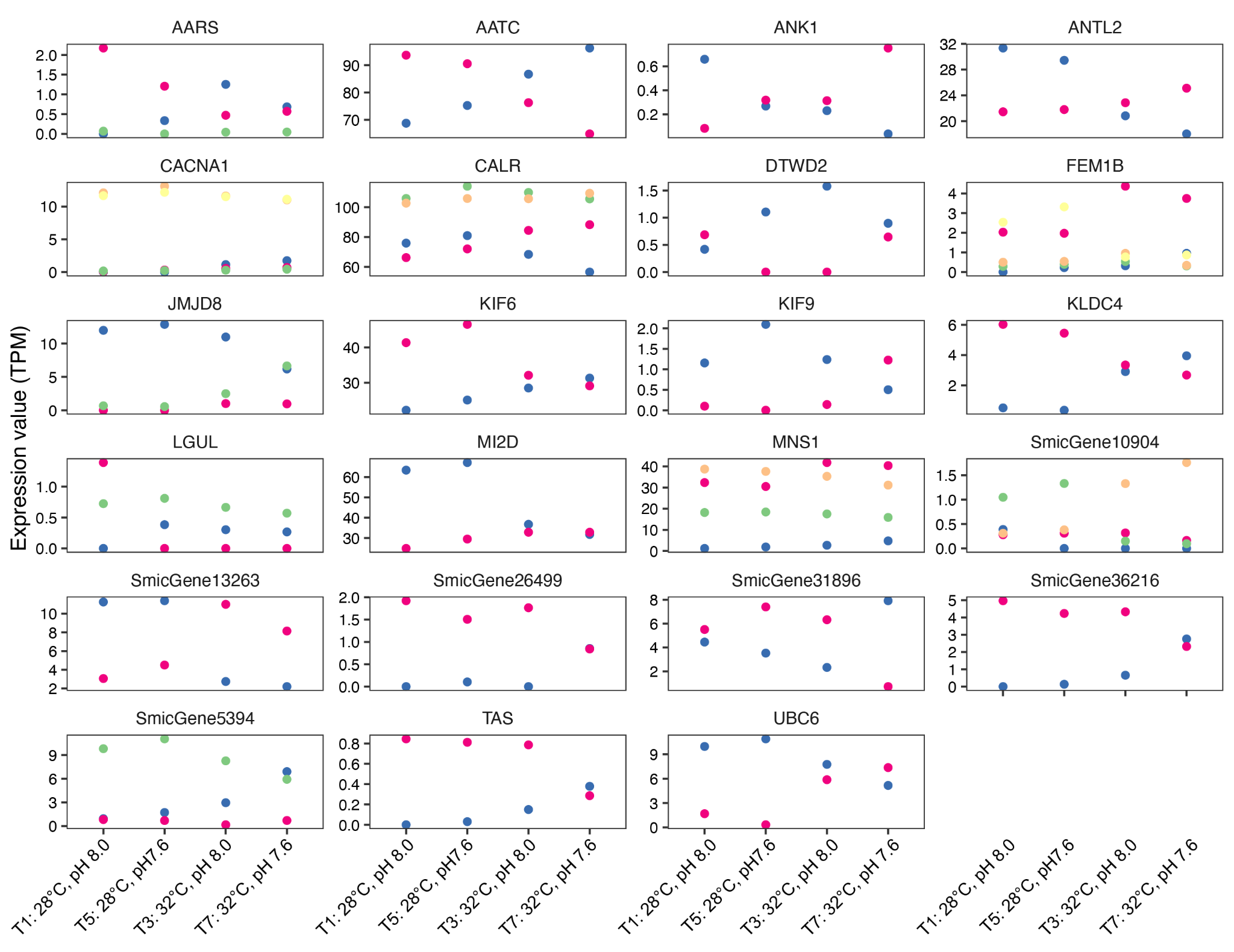
**Figure S2.** Genes that exhibit isoform switching across treatments. Differently colored dots represent different isoforms.

**Supplementary Tables**

**Table S1.** Seawater parameters in the experimental treatments. Average values (± standard error) during the sustained exposure period (days 11-18). TA and DIC were measured on days 13 and 16, except for T4 and T8, which were measured only on day 13. Temp, temperature; TA, total alkalinity; DIC, dissolved inorganic carbon, *p*CO_2_, partial pressure of carbon dioxide; HCO_3_^-^, bicarbonate; CO_3_^-2^, carbonate; Ωarag, aragonite saturation state; Ωcal, calcite saturation state; Sal, salinity; DO, dissolved oxygen.

| Treatment  (°C/pH) | Temp (°C) | pH | TA (µmol/kg) | DIC (µmol/kg) | *p*CO_2_ (µatm) | HCO_3_^-^  (µmol/kg) | CO_3_ ^-2^  (µmol/kg) | Ωarag | Ωcal | Sal  (o/oo) | DO  (mg/L) |
| --- | --- | --- | --- | --- | --- | --- | --- | --- | --- | --- | --- |
| T1: 28°C/8.0 | 28.48  ±0.03 | 8.06  ±0.01 | 2177.48  ±24.03 | 2082.21  ±19.43 | 951.26  ±108.67 | 1910.99  ±41.51 | 108.82  ±9.54 | 1.77  ±0.16 | 2.70  ±0.24 | 30.91  ±0.06 | 5.03  ±0.04 |
| T2: 30°C/8.0 | 30.36  ±0.05 | 8.02  ±0.01 | 2221.60  ±3.46 | 2117.55  ±20.66 | 1104.19  ±115.52 | 1978.31  ±25.84 | 99.77  ±9.26 | 1.62  ±0.15 | 2.47  ±0.23 | 31.23  ±0.02 | 4.92  ±0.03 |
| T3: 32°C/8.0 | 32.34  ±0.07 | 8.03  ±0.01 | 2220.84  ±5.19 | 2096.09  ±20.52 | 965.58  ±104.99 | 1952.36  ±27.10 | 110.12  ±9.16 | 1.79  ±0.15 | 2.73  ±0.23 | 31.15  ±0.03 | 4.82  ±0.03 |
| T4:  34°C/8.0 | 34.09  ±0.11 | 7.98  ±0.01 | 2222.80  ±2.14 | 2068.55  ±8.66 | 775.33  ±30.43 | 1919.15  ±12.76 | 124.69  ±4.50 | 2.04  ±0.07 | 3.10  ±0.11 | 31.07  ±0.11 | 4.48  ±0.10 |
| T5:  28°C/7.6 | 28.49  ±0.04 | 7.65  ±0.01 | 22221.10  ±3.09 | 2202.15  ±8.50 | 1912.03  ±101.24 | 2077.15  ±9.23 | 59.09  ±2.95 | 0.96  ±0.05 | 1.47  ±0.08 | 30.93  ±0.05 | 5.12  ±0.03 |
| T6:  30°C/7.6 | 30.41  ±0.06 | 7.62  ±0.01 | 2231.33  ±4.74 | 2279.49  ±57.20 | 2169.46  ±177.73 | 2096.02  ±13.29 | 55.62  ±4.02 | 0.91  ±0.07 | 1.38  ±010 | 30.89  ±0.35 | 4.84  ±0.04 |
| T7:  32°C/7.6 | 32.40  ±0.06 | 7.62  ±0.01 | 2240.09  ±9.11 | 2229.11  ±17.06 | 2025.00  ±141.66 | 2100.71  ±16.85 | 57.30  ±3.65 | 0.93  ±0.06 | 1.42  ±0.09 | 31.18  ±0.03 | 4.86  ±0.04 |
| T8:  34°C/7.6 | 33.98  ±0.09 | 7.59  ±0.01 | 2217.68  ±3.65 | 2203.08  ±9.24 | 1901.30  ±133.24 | 2074.28  ±8.21 | 58.91  ±3.34 | 0.96  ±0.06 | 1.46  ±0.08 | 31.33  ±0.10 | 4.39  ±0.11 |

**Table S2.** Mean Symbiodiniaceae densities under different temperature and pH treatments.

| **Treatment (°C/pH)** | **n** | **Cells per gram wet weight**  **(mean ± standard error)** |
| --- | --- | --- |
| T1: 28**°**C / pH 8.0 | 8 | 2.05 ± 0.32 x 10^8^ |
| T5: 28**°**C / pH 7.6 | 8 | 2.06 ± 0.38 x 10^8^ |
| T2: 30**°**C / pH 8.0 | 8 | 1.46 ± 0.54 x 10^8^ |
| T6: 30**°**C / pH 7.6 | 8 | 1.58 ± 0.07 x 10^8^ |
| T3: 32**°**C / pH 8.0 | 8 | 5.78 ± 1.57 x 10^7^ |
| T7: 32**°**C / pH 7.6 | 8 | 6.65 ± 0.99 x 10^7^ |

**Table S3.** *Tridacna gigas* *de novo* transcriptome assembly statistics.

| ***De novo* assembly** | |
| --- | --- |
| Transcripts | 207245 |
| Genes | 180246 |
| GC% | 44.71 |
| N50 length (bp) | 1390 |
| Ex90N50 length (bp) | 1635 |
| Protein-coding transcripts | 77228 |
| Host transcripts (metazoa hits) | 19842 |
| Symbiont transcripts (alveolate hits) | 34328 |
| No hits to nr | 23058 |
| **Non-redundant protein-coding transcriptome** | |
| Transcripts | 54170 |
| Genes | 46296 |
| GC% | 48.98 |
| N50 length (bp) | 2203 |
| Ex90N50 length (bp) | 1921 |
| ***Host bin*** |  |
| Transcripts | 19842 |
| Genes | 16837 |
| GC% | 40.72 |
| N50 length (bp) | 2603 |
| ***Symbiont bin*** |  |
| Transcripts | 34328 |
| Genes | 29459 |
| GC% | 54.53 |
| N50 length (bp) | 2010 |
| **BUSCO** | |
| Host vs mollusca_odb12 | C: 86.7% [S: 78.0%, D: 8.7%], F: 6.4%, M: 6.8%, n: 4421 |
| Host vs metazoa_odb12 | C: 95.5% [S: 87.5%, D: 8.0%], F: 2.8%, M: 1.6%, n: 672 |
| Host vs alveolata_odb12 | C: 87.9% [S: 82.8%, D: 5.1%], F: 8.1%, M: 4.0%, n: 99 |
| Host vs bacteria_odb12 | C: 31.9% [S: 28.4%, D: 3.4%], F: 8.6%, M: 59.5%, n: 116 |
| Symbiont vs mollusca_odb12 | C: 12.1% [S: 9.3%, D: 2.7%], F: 3.3%, M: 84.6%, n: 4421 |
| Symbiont vs metazoa_odb12 | C: 34.2% [S: 30.4%, D: 3.9%], F: 13.5%, M: 52.2%, n: 672 |
| Symbiont vs alveolata_odb12 | C: 94.9% [S: 84.8%, D: 10.1%], F: 4.0%, M: 1.0%, n: 99 |
| Symbiont vs bacteria_odb12 | C: 50.0%, [S: 31.9%, D: 18.1%], F: 6.0%, M: 44.0%, n: 116 |

**Table S4.** Summary of best BLAST hits to Symbiodiniaceae genes, showing species level assignments, number of genes, and relative abundance (%).

| **Species** | **Number of genes with best hit** | **Relative abundance (%)** |
| --- | --- | --- |
| *Cladocopium infistulum* | 9677 | 32.85 |
| *Cladocopium proliferum* | 5984 | 20.31 |
| *Cladocopium* sp. (Y103 isolate) | 7612 | 25.84 |
| *Cladocopium* sp. (C15 ITS2-subtype) | 3413 | 11.59 |
| *Breviolum minutum* | 1430 | 4.85 |
| *Fugacium kawagutii* | 606 | 2.06 |
| *Durusdinium trenchii* | 430 | 1.46 |
| *Symbiodinium microadriaticum* | 296 | 1.00 |
| Unassigned | 11 | 0.04 |

**Supplementary datasets**

**Dataset 1.** Daily mortality and survival rates

**Dataset 2.** Differentially expressed genes

**Dataset 3.** Functional enrichment analysis

**Dataset 4.** Semantic similarity clustering of enriched functions

**Dataset 5.** Protein interaction network from STRING

**Dataset 6.** Isoform switching
